# The latent geometry of the human protein interaction network

**DOI:** 10.1101/213165

**Authors:** Gregorio Alanis-Lobato, Pablo Mier, Miguel A. Andrade-Navarro

## Abstract

To mine valuable information from the complex architecture of the human protein interaction network (hPIN), we require models able to describe its growth and dynamics accurately. Here, we present evidence that uncovering the latent geometry of the hPIN can ease challenging problems in systems biology. We embedded the hPIN to hyperbolic space, whose geometric properties reflect the characteristic scale invariance and strong clustering of the network. Interestingly, the inferred hyperbolic coordinates of nodes capture biologically relevant features, like protein age, function and cellular localisation. We also realised that the shorter the distance between two proteins in the embedding space, the higher their connection probability, which resulted in the prediction of plausible protein interactions. Finally, we observed that proteins can efficiently communicate with each other via a greedy routeing process, guided by the latent geometry of the hPIN. When analysed from the appropriate biological context, these efficient communication channels can be used to determine the core members of signal transduction pathways and to study how system perturbations impact their efficiency.

## Introduction

Proteins are very complex machines in and of themselves, but their interactions with other proteins foster the formation of a very intricate molecular system. This level of complexity has propelled the development of methods to facilitate the analysis of protein interaction networks^1^ and has led to notable advances in biology and medicine^2–6^.

Of special interest are a series of algorithms and models that advocate for the existence of a geometry underlying the structure of complex networks, shaping their topology^7–14^(we refer the reader to^15^ for an extensive review on the subject). In particular, the Popularity-Similarity model (PSM) sustains that the emergence of strong clustering and scale invariance, properties common to most complex networks, is the result of certain trade-offs between node popularity and similarity^13^. This model has a geometric interpretation in hyperbolic space (ℍ^2^), where distance-dependent connection probabilities lead to link formation, accurately describing the growth of complex systems^12,13,16–19^.

In the PSM, the *N* nodes comprising a network lie within a circle of radius *R* ~ ln *N*, at polar coordinates (*r_i_*,*θ_i_*). The radial coordinate *r_i_* represents the popularity or seniority status of a node *i* in the system. Nodes that joined the system first have had more time to accumulate links and are close to the circle’s centre, whereas younger nodes lie on the circle’s periphery and have only a few partners. The angular coordinate *θ_i_* allows one to determine how similar a node *i* is to others. Finally, the hyperbolic distance between nodes, *d_ℍ^2^_* (*s*, *t*) ≈ *r_s_* + *r_t_* + 2 ln (*θ_st_*/2), abstracts the optimisation process mentioned above, in which a new node aims at forming a tie not only with the most popular system components but also with the ones that are most similar to it^13^.

The PSM is markedly appealing to network biologists because the human protein interaction network (hPIN), the focus of this study, exhibits an approximately scale-free node degree distribution and has a strong clustering (see Table S1). Furthermore, uncovering the hidden geometry of the hPIN could ease challenging problems in systems biology^20^, allowing us to address them from a geometric perspective. For example, the prediction of protein interactions would translate into the identification of disconnected protein pairs that are unexpectedly close to each other in the network’s latent space.

To investigate whether ℍ^2^ represents a good host space for the hPIN, we developed an accurate and efficient algorithm for hyperbolic network embedding^21^ and explored whether the popularity and similarity dimensions inferred for each protein have a biological interpretation. Furthermore, we exploited the hyperbolic distance between proteins for link prediction and the reconstruction of signal transduction pathways.

## Results and discussion

### Uncovering the latent geometry of the human protein interactome

We constructed a protein network with high-quality interactions from the Human Integrated Protein-Protein Interaction rEference or HIPPIE^22,23^, which reports interactions with a confidence score that reflects their experimental evidence (see Methods and Supplementary Data S1). The resulting network was embedded to the two-dimensional hyperbolic plane ℍ^2^ using LaBNE+HM^21^, a method that combines manifold learning^24^ and maximum likelihood estimation^25^ to uncover the hidden geometry of complex networks (see Methods). Once the hyperbolic coordinates of each protein in the network were inferred (see Supplementary Data S2), we proceeded to analyse whether these coordinates are meaningful or not from a biological point of view.

### Radial coordinates and protein evolution

As discussed in the introduction, the popularity component of the PSM (radial coordinates of nodes in ℍ^2^) is associated with the seniority status of network nodes. Old nodes tend to be close to the centre of the hyperbolic circle containing the system and their degree (their number of direct neighbours) tends to be much higher than that of young peripheral nodes^26^. To verify if our mapping reflects these tendencies, we assigned proteins to six different age groups according to the existence of evolutionarily-related counterparts in other organisms (see Fig. 1a, Methods and Supplementary Data S2).

**Figure 1:**
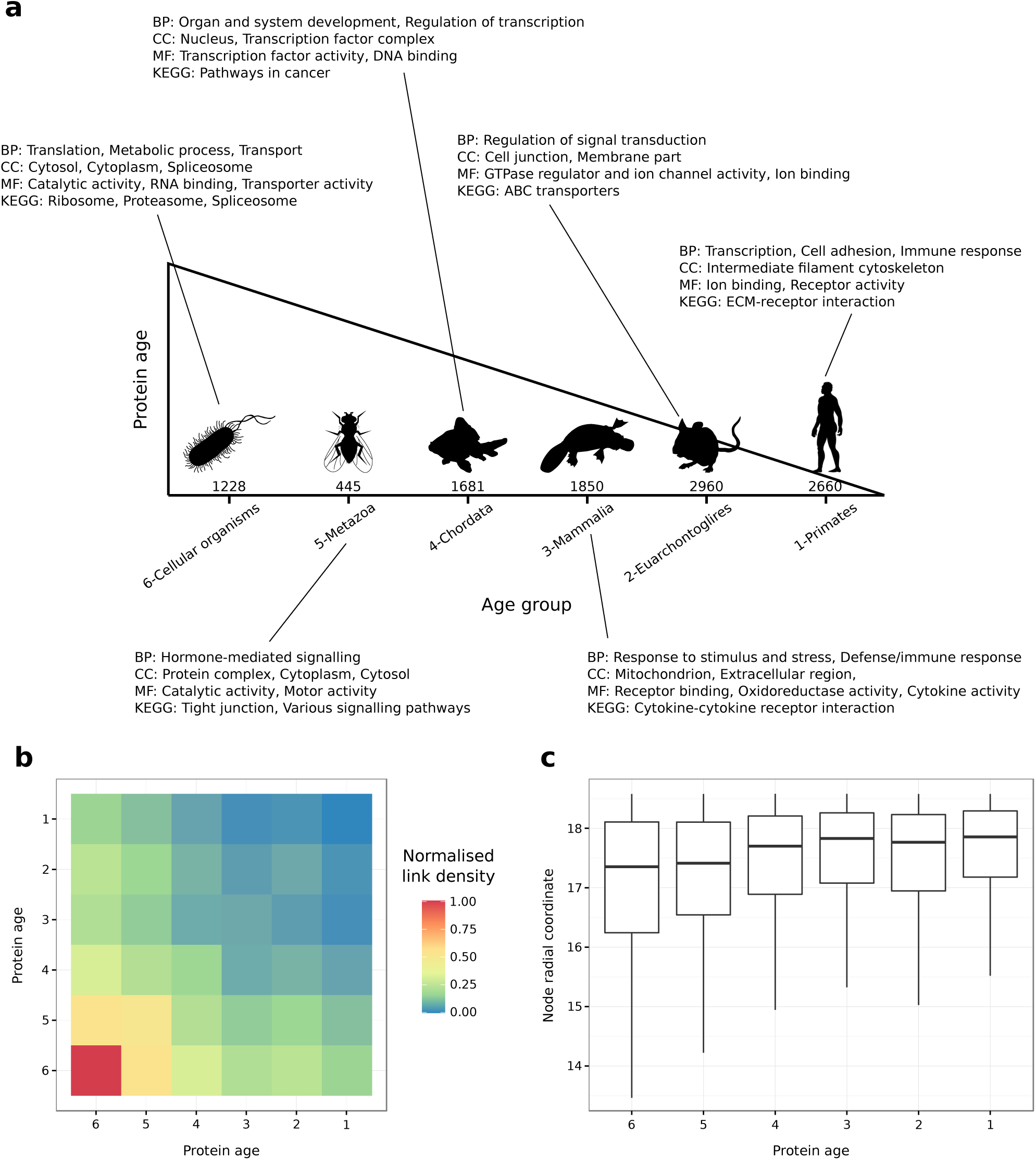
Popularity and protein evolution. (**a**) Proteins in the constructed hPIN were clustered into six different age groups (the number of proteins in each one is indicated). The over-represented biological functions and compartments in each group were determined via Gene Ontology and KEGG pathway enrichment analyses (BP: Biological Process, CC: Cellular Compartment, MF: Molecular Function, KEGG: Kyoto Encyclopedia of Genes and Genomes). (**b**) The normalised link density within and between age groups. (**c**) The distribution of inferred radial coordinates for the proteins in each age group.

The validity of the determined age groups is supported by the node degree and biological functions of their constituent proteins. While old nodes have high degrees and are involved in essential functions, like metabolic processes or protein translation, younger nodes have only a few direct partners and are in charge of more specialised processes, like organ development and immune response (see Fig. 1a, S1a and Supplementary Data S3). Moreover, there is a strong link density (observed links between proteins relative to the total number of possible links that can occur) within and between old age groups, which is reduced within and between the young ones (see Fig. 1b and Methods). This is in agreement with previous observations that there is a core of old highly interconnected proteins, surrounded by younger proteins with no interactions between them but dependent on the old core^27,28^. All these results cannot be replicated if protein ages are assigned randomly (see Figs. S1b,c).

Finally, we checked the inferred radial coordinates of the proteins in each group and, in agreement with the PSM, old proteins are closer to the centre of the hyperbolic circle compared to younger ones (see Fig. 1c). The observed trend is an indication that the radial positions of proteins in ℍ^2^ encode information about their evolutionary origin.

### Angular coordinates and the spatio-functional organisation of the cell

The similarity component of the PSM (angular coordinates of nodes in ℍ^2^) abstracts the characteristics that make a node similar to others. To investigate the biological meaning of this dimension, we computed the difference between consecutive protein angles to identify big gaps separating agglomerations of similar proteins. To determine cluster membership, we chose the gap size *g* that produced clusters with at least 10 proteins. This choice allowed us to perform meaningful functional enrichment analyses of each group (see Fig. S2 and Methods).

As shown in Fig. 2a, the similarity dimension captures the functional and spatial organisation of the cell, and this is supported by the three aspects of the Gene Ontology and by KEGG Pathways (see Supplementary Data S4 and Fig. S3). For example, the over-represented biological process of cluster 8 is *transcription.* The cellular compartment where this process takes place, the *nucleus*, is also enriched, as well as the molecular functions *DNA binding* and *transcription factor activity* together with the *basal transcription factors* pathway.

**Figure 2:**
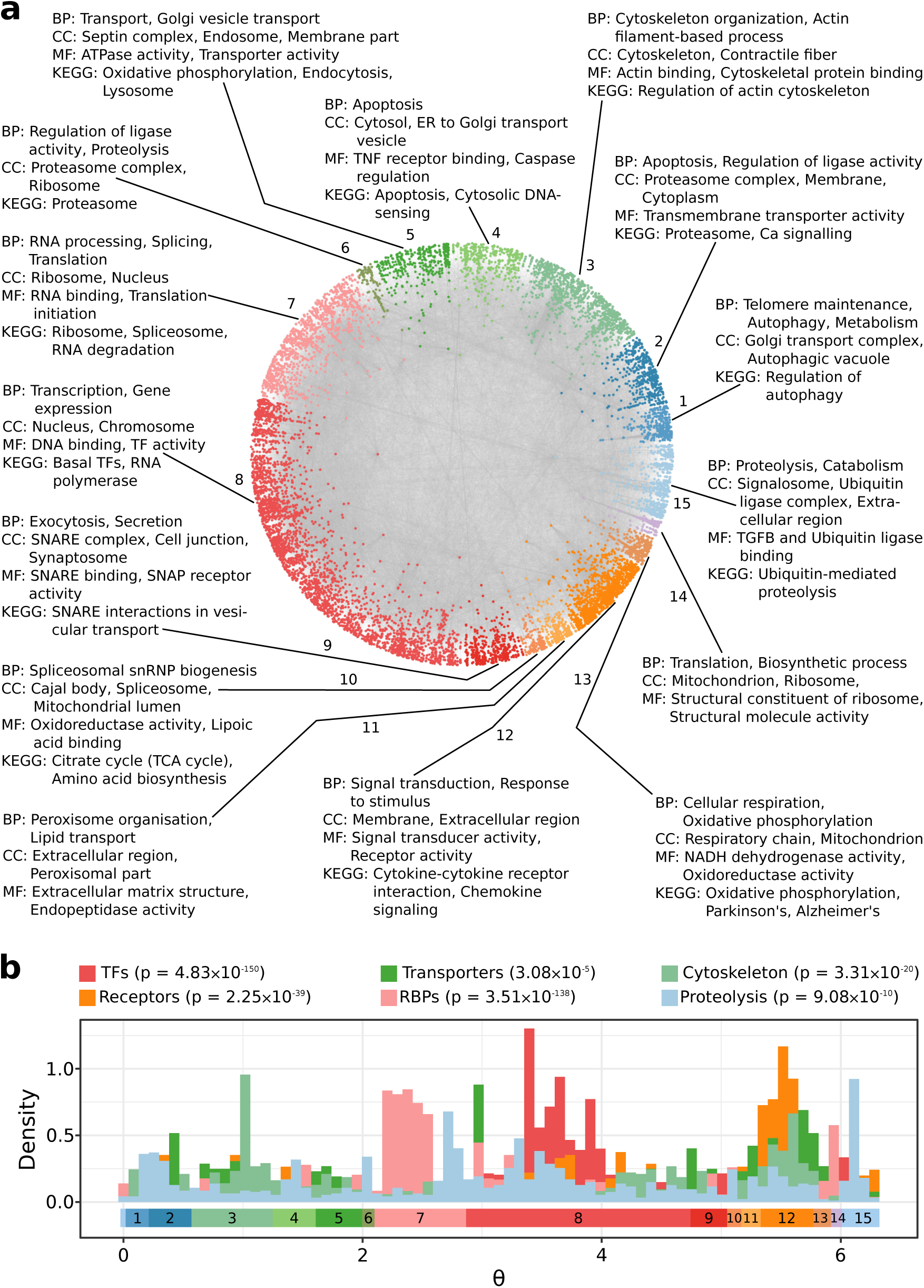
Similarity and protein function. (**a**) Protein clusters identified by big gaps separating groups of proteins in the angular dimension of hyperbolic space. The over-represented biological functions and compartments in each cluster were determined via Gene Ontology and KEGG pathway enrichment analyses (BP: Biological Process, CC: Cellular Compartment, MF: Molecular Function, KEGG: Kyoto Encyclopedia of Genes and Genomes). Each cluster has been assigned a numeric identifier (1-15). (**b**) The distribution of inferred angular coordinates for proteins with specific molecular functions (TFs: Transcription Factors, RBPs: RNA-binding proteins). The reported p-values (adjusted with the Benjamini-Hochberg method), which are the result of a Fisher’s exact test, highlight that these protein classes agglomerate non-randomly within their corresponding similarity-based cluster from a. The start and end of each one of these clusters are indicated across the [0, 2*π*] range, below the histograms.

We also consulted various databases, censuses and atlases to determine whether proteins in the hPIN act as transcription factors, receptors, transporters, RNA-binding proteins, constituents of the cytoskeleton or in proteolysis/ubiquitination processes (see Methods and Supplementary Data S2). Fig. 2b shows the distribution of inferred angles for these protein classes and highlights how they agglomerate in the similarity-based clusters enriched for their particular activity, in numbers that are significantly higher than expected by chance (see Methods). For example, RNA-binding proteins accumulate in cluster 7, which, as expected, is enriched for RNA processing and protein translation. Also, nodes involved in marking proteins with ubiquitin for their degradation via the proteasome, though more dispersed across the full angular dimension, are more common in the clusters enriched for ubiquitination and proteolysis (1, 2, and 15).

To study whether the clusters suggested by the angular coordinates of proteins could have been detected with a traditional community detection method, we applied the Louvain algorithm to the hPIN ^29^. This method identified communities that do not correspond with the obtained similarity-based clusters (see Figs. S4a-d).The Louvain-based communities are either enriched for very specific biological processes or not enriched for any process in particular (see Supplementary Data S5). This outcome suggests that they represent protein complexes or groups of a few proteins that, together, play roles in very particular functions (see Fig. S4d). In contrast, the angular clusters are formed by proteins with roles in more general pathways (see Supplementary Data S4) that can be analysed in more detail if smaller gaps between angles are considered (see Figs. S2, S4c and Methods).

The results presented so far correspond to an hPIN formed by interactions with HIPPIE confidence scores ≥ 0.72 (see Methods), which means that they are well-supported by experimental evidence. However, this also means that the considered interactome is vastly incomplete. To test if our findings are robust to network topology changes (e.g. higher presence of false negatives if a more stringent score is used or more false positives if the score is less conservative), we constructed hPINs with varying quality levels (see Table S1). Fig. S5 shows that regardless of the assessed confidence score, the inferred protein coordinates lead to the same conclusions: old proteins tend to be closer to the centre of ℍ^2^ than young ones^27,28^and proteins with specific molecular functions cluster together in the angular dimension. We expect these observations to hold true or even improve, as hPIN charting efforts enhance network coverage and reliability^5,6^.

### Hyperbolic distances and protein interaction prediction

Now that the two dimensions of the PSM have been interpreted in a biological context, we can use them to compute hyperbolic distances between proteins. Fig. 3a shows connection probabilities (the fraction of connected node pairs, amongst all pairs separated by a certain distance) as a function of the hyperbolic separation between proteins. In concordance with what the PSM predicts for a network with the same structural characteristics as the hPIN, we can see that, according to the coordinates inferred with LaBNE+HM, if two proteins are very close to each other, they most certainly interact. On the other hand, if proteins are far apart, their probability of interaction is very low. Additionally, protein interactions with high HIPPIE confidence scores are closer to each other than proteins with low scores (see Fig. S6).

**Figure 3:**
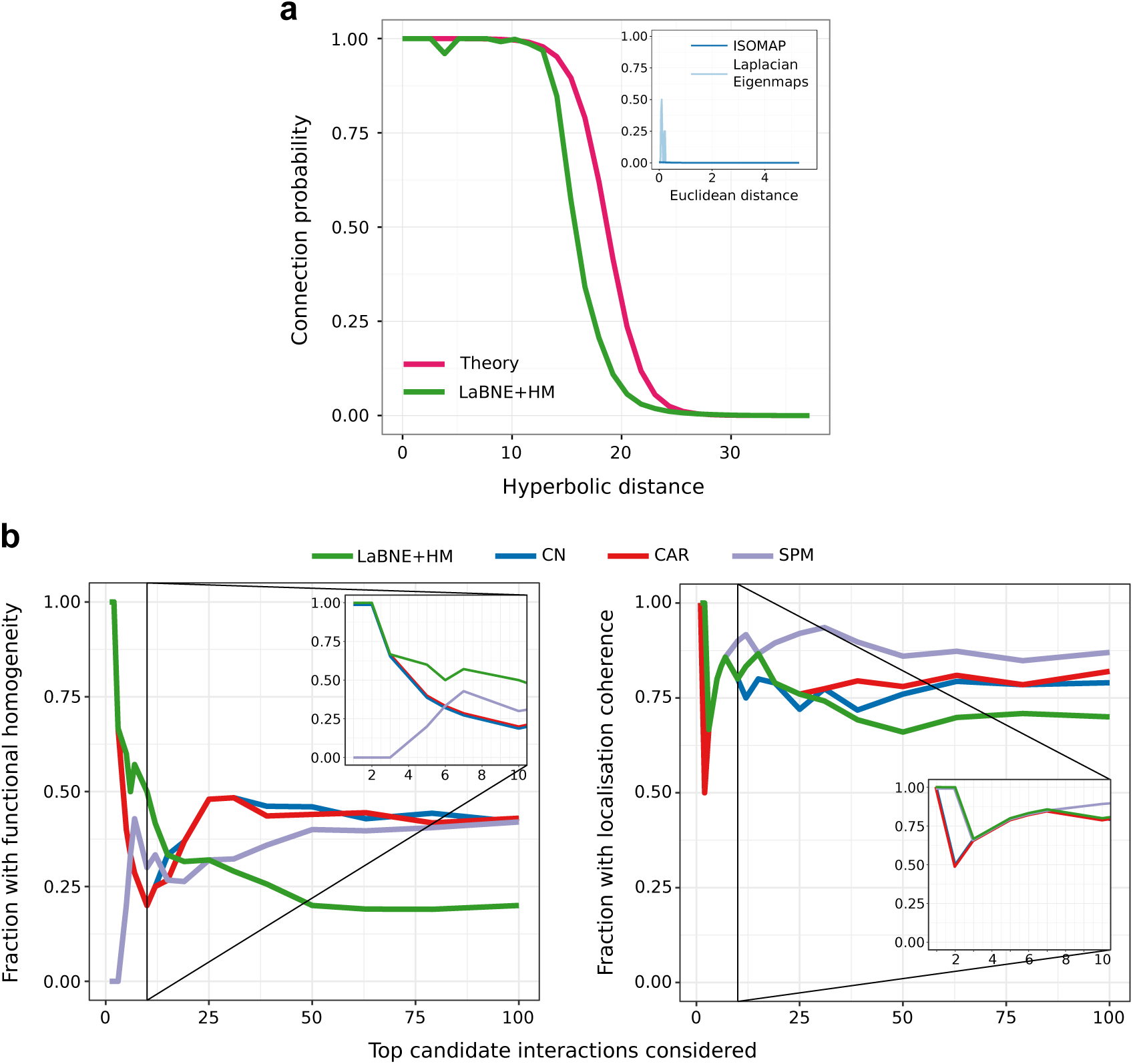
Hyperbolic distance and likelihood of interaction. (**a**) Connection probability as a function of the hyperbolic or Euclidean (inset) separation between protein pairs. The probabilities predicted by the Popularity-Similarity model (Theory) and the ones obtained by mapping the network to a geometric space with LaBNE+HM, ISOMAP and Laplacian Eigenmaps are shown. (**b**) We compared the top-100 disconnected proteins that are closest to each other in ℍ^2^ (LaBNE+HM) with candidate protein interactions from representative link predictors of different classes (see Fig. S7 for the complete analysis). The plot shows how the fraction of potential interactions with functional homogeneity and localisation coherence changes as more protein pairs are assessed. The insets focus on the top-10 candidate pairs. CN: Common Neighbours, CAR: Cannistraci-Alanis-Ravasi index, SPM: Structural Perturbation Method.

We tried to replicate the above findings by embedding the hPIN into the two-dimensional Euclidean space, using two different techniques^30,31^(we refer the reader to^9,10,14^for details on how these network embeddings are performed). The resulting connection probabilities are far from what the mapping to ℍ^2^ achieves (see inset in Fig. 3a), further endorsing the suitability of this space to describe complex networks like the hPIN.

These results encouraged us to check whether the 100 hyperbolically-closest disconnected protein pairs represent plausible protein interactions. Fig. 3b shows that LaBNE+HM’s predictions are more biologically meaningful than those from representatives of different link prediction classes^32,33^(see Fig. S7 for the complete analysis, as well as the Methods and Supplementary Data S6), especially if we focus on the top-10 candidates: non-adjacent proteins that are close in ℍ^2^ play roles in at least one common pathway (functional homogeneity) and localise to the same cellular compartments (localisation coherence).

Our top prediction, for example, involves proteins SUMO2 and p65 and is supported by recent studies in mouse and human. After observing that over-expression of SUMO2 derives in the lack of nuclear p65, a group working with mouse dendritic cells proposed that SUMO2 traps p65 in the cytoplasm and avoids its translocation to the nucleus^34^. Further supporting this hypothesis, Liu and colleagues observed that the transfection of human hepatocarcinoma with increasing doses of SUMO2 gradually increases cytoplasmic p65 levels, whereas knock-down of SUMO2 decreases them^35^.

The second-top prediction implicates proteins MYC and GPC1. Qiao and colleagues found that the ectopic expression of GPC1 can significantly induce both the MYC protein and its transcription in a cell type-independent manner^36^. On the other hand, inhibition of GPC1-induced MYC transactivation almost completely overturns DNA re-replication, suggesting an important role of both proteins in the mediation of cellular effects and signalling ^36^.

Although the other link prediction approaches improve as more candidates are evaluated, we cannot discard that some of LaBNE+HM’s predictions are actually part of the same pathway or organelle, as pathway membership and protein localisation references are still incomplete. A sign of this lack of annotations is that only ~ 20% of the top-100 potential interactions identified by each prediction method are reported in HIPPIE v2.0 (see Fig. S8a) and an average of around three was confirmed by a recent large-scale network charting effort^6^ (see Fig. S8b). This means that there is no experimental evidence for the interaction of most these protein pairs, a problem that proteome-scale and unbiased protein network mapping endeavours are addressing^5^.

### Greedy routeing and signal transduction

Hyperbolic distances can also be used to study signal transduction pathways, the way in which cells communicate with each other and respond to environmental changes^37^. These pathways normally start with a signal stimulating a cell membrane receptor, which leads to the activation of a series of proteins, until the signal reaches the nucleus, where a transcription factor binds DNA and transcribes target genes^38^.

Interestingly, signals travel from source to target with the former not having knowledge of the global protein network structure^8,12^. Proteins can only activate or repress their direct neighbours in the hPIN, and these stimuli cascade through the network in the same way, until the end of the pathway^38^. This prompted us to investigate whether a signal can effectively reach its target, using the shortest possible path, via a greedy routeing process. In greedy routeing, the inferred hyperbolic coordinates of proteins are used as addresses that allow a signal to get closer and closer to the target. The routeing process starts with the source checking which one of its direct neighbours is hyperbolically closest to the target and sends the signal there. This new node checks amongst its direct partners for the one closest to the target, and so on, until the signal reaches its destination. If in the delivery process, a neighbour sends the signal to the previously visited protein, i.e. it falls into a loop, the signal is dropped and the delivery is flagged as unsuccessful^12^.

We sent signals from 1000 randomly selected sources to 1000 randomly selected targets, computed the percentage of successfully delivered signals and repeated the experiment 100 times. Fig. 4a shows the average efficiencies. Note that if signals travel to the neighbour that is radially or angularly closest to the target, greedy routeing is not as efficient as when the hyperbolic distances are used, underlining the importance of both dimensions for the proper navigation of the hPIN^12,21^. Moreover, the hop stretch (greedy path length divided by shortest path length) is close to 1 (see Fig. 4b), which means that greedy paths, guided by the network’s latent geometry, are very often shortest paths.

**Figure 4:**
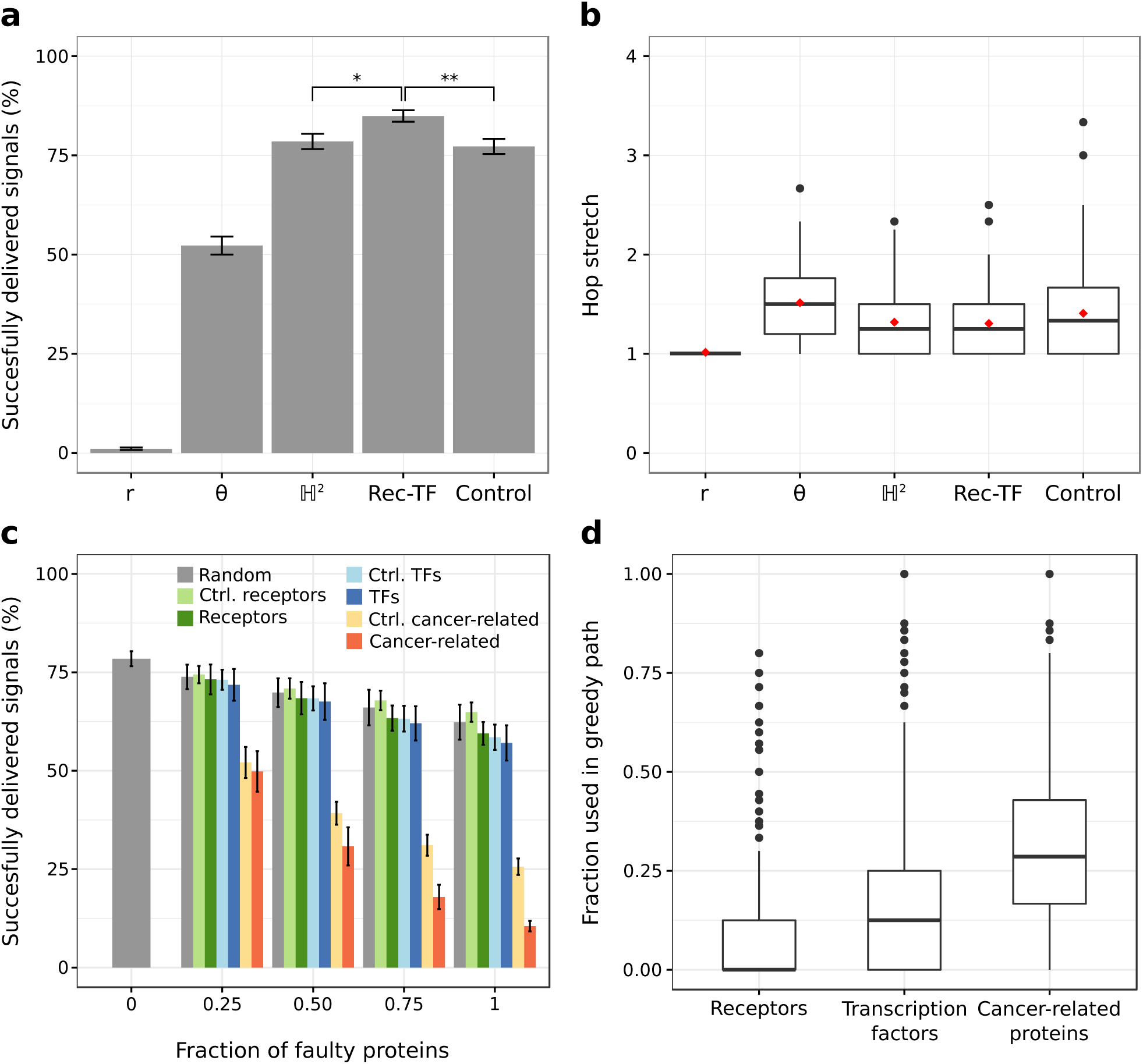
Navigability of the human protein network. (**a**) Percentage of successfully delivered signals travelling from random sources to random targets (using the neighbour radially *r*, angularly *θ* or hyperbolically ℍ^2^ closest to the target), from receptors to transcription factors (Rec-TF) or from proteins that are neither receptors nor transcription factors, but with degrees similar to their counterparts (Control). \**p* = 1.898 × 10^−34^, \*\**p* = 1.233 × 10^−34^, Mann-Whitney U test. (**b**) Hop stretches for all the cases presented in a. Average hop stretches are reported with red diamonds. (**c**) Percentage of successfully delivered signals when increasing levels of faulty proteins are introduced. Faulty proteins are chosen at random or from a pool with the same number of receptors, transcription factors, cancer-related proteins, control receptors (Ctrl. receptors), control transcription factors (Ctrl. TFs) or control cancer-related proteins (Ctrl. cancer-related). See a for details on the controls. (**d**) Distribution of the fraction of receptors, transcription factors and cancer-related proteins used in 1000 different greedy paths. Error bars correspond to standard deviations.

Given the biological importance of signal transduction, we hypothesised that it should be more efficient to send signals from receptors to transcription factors, and that is indeed the case (*p* = 1.898 × 10^−34^, see Fig. 4a). The difference is also significant with respect to the use of proteins that are neither receptors nor transcription factors, but that have degrees similar to their counterparts (*p* = 1.233 × 10^−34^, see Fig. 4a and Fig. S9a,b). Here, we refer to them as control receptors and control transcription factors, respectively.

We also explored the effects of defective proteins in greedy routeing efficiency. A faulty protein drops any signal it receives, making routeing unsuccessful. From a biological perspective, unsuccessful routeing could be modelling cellular mechanisms compensating the effects caused by mutations or insufficient protein levels. In some situations, these defects could manifest as disease phenotypes.

As depicted in Fig. 4c, the increasing introduction of defective receptors or transcription factors impacts greedy routeing efficiency more than the introduction of faulty proteins at random or from the pool of control receptors or control transcription factors. We tested this using pools with the same amount of receptors and transcription factors to make sure that the observed effects were not due to different abundances of these protein types in the hPIN. Interestingly, faulty nodes from a pool of cancer-related proteins (see Methods) severely affect network navigability compared to transcription factors, receptors and even control cancer proteins (see Fig. 4c and Fig. S9c). This impact on routeing efficiency cannot be attributed to cancer proteins having more connections, as their degree distribution is similar to that of transcription factors and receptors (see Fig. S9). Rather, it could be explained by how often cancer-related proteins are part of greedy paths (see Fig. 4d) and motivates a deeper investigation of the relationship between network navigation, function and disease, which is outside the scope of this work.

### Geometry-based pathway reconstruction

One of the major challenges in systems biology is the determination of the chain of reactions that guides signals from receptors in the cell membrane to transcription factors in the nucleus^39^. Although current experimental technologies enable the identification of the proteins in charge of sensing the cell’s environment and the deduction of the downstream effects of these sensory inputs, building the complete set of interactions that are part of signalling pathways still requires extensive and time-consuming manual curation efforts^39,40^. As a result, the development of automatic pathway reconstruction methods is a field of active research^39–43^. Such methods aim at establishing pathway members and their interactions, given only two anchoring points: the receptor or source of the pathway and the target transcriptional regulator^39^.

We explored the extent to which well-established signal transduction pathways can be recapitulated by navigating the latent geometry of the hPIN with greedy routeing. Note that our goal was not the full reconstruction of pathways, with all their diversions, loops and buffering controls. Rather, our objective was to study whether the inferred network geometry can guide signals through the core pathway members.

Using greedy routeing and traditional shortest paths, we sent signals from canonical sources to canonical transcriptional regulators of the 24 signal transduction pathways listed in KEGG^44^ (see Supplementary Data S7). Then, we computed the fraction of proteins that are part of the resulting greedy/shortest paths and that are reported pathway members in a dataset integrating KEGG ^44^, Reactome^45^ and WikiPathways^46^ information (see Methods). Figs. 5a and S10 show that, in 70% of the cases, greedy paths are as good as or better than shortest paths because they contain more proteins that are in fact part of the analysed pathway. Along with this, hop stretches fluctuate around 1, indicating that the navigated greedy paths, found using local information only, are often shortest paths. However, longer greedy paths with just a small fraction of reported pathway members are also interesting, as they may contain new pieces of the signal transduction machinery. In the following paragraphs, we discuss a few individual cases from Fig. 5a, including examples of how this framework can be exploited to generate compelling biological hypotheses.

**Figure 5:**
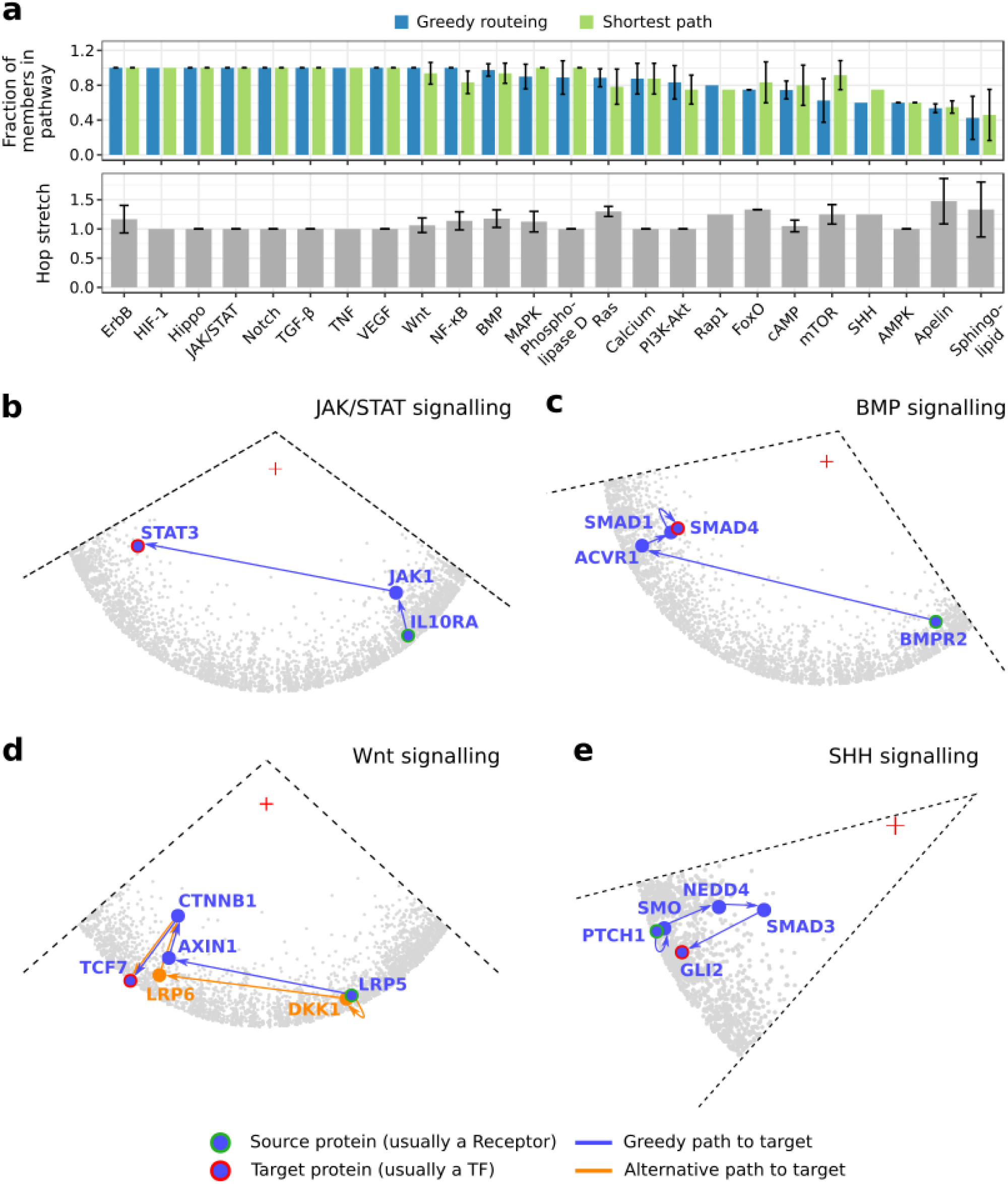
Geometry-based pathway reconstruction. (**a**) Signals were routed from receptors to transcriptional regulators of the 24 signal transduction pathways listed in the Kyoto Encyclopedia of Genes and Genomes (KEGG). Greedy routeing and shortest-paths were employed. The fraction of greedy path and shortest path members that are part of each pathway is reported, together with the hop stretch (greedy path length divided by shortest path length). When more than one source or target was considered, the average fraction is reported. Error bars correspond to standard deviations. Reconstruction of the (**b**) JAK/STAT, (**c**) BMP, (**d**) Wnt and (**e**) SHH signal transduction pathways by navigation of the latent geometry of the hPIN with greedy routeing. A red cross indicates the centre of the hyperbolic circle containing the hPIN. In the case of **d**, a signal can successfully reach TCF7, via an alternative greedy path, when AXIN1 is removed from the network (*in silico* knock-out experiment).

Activation of the JAK/STAT pathway stimulates cell proliferation, differentiation, migration and apoptosis^47^. The key elements of this pathway, the STAT transcription factors, are inactive in unstimulated cells and reside in the cytoplasm. When cytokine membrane receptors are activated, JAK protein kinases, bound to such receptors, phosphorylate STAT proteins, which then translocate to the nucleus, where they trigger the transcription of target genes^38,47^. It turns out that if we navigate the latent geometry of the hPIN from a cytokine receptor (IL10RA) to a STAT transcription factor (STAT3), the described series of signalling events can be recapitulated (see Fig. 5b).

Analysis of the BMP pathway, crucial for bone and cartilage development^48^, leads to a similar result. In this pathway, BMP signals bind to a complex of BMP and activin receptors. Once activated, these receptors phosphorylate SMAD1, SMAD5 or SMAD8 proteins, which then form a complex with SMAD4, the transcription factor that translocates to the nucleus for gene regulation^48^. Fig. 5c shows how the geometry of the hPIN guides a signal from the BMP receptors (BMPR2 and ACVR1) to the SMAD1 protein, to finally reach SMAD4.

Wnt signalling, a well-characterised pathway with an important role in embryonic development^49^, can also be reconstructed with greedy routeing (see Fig. 5d). In its canonical form, this pathway is activated when a Wnt signal stimulates the LRP membrane receptors (LRP5 and LRP6), leading to their association with a multiprotein complex containing AXIN1. This event stabilises the *β*-catenin protein (CTNNB1), trapped by the multiprotein complex before the pathway activation, and allows its translocation to the nucleus, where it binds transcription factor TCF7 and induces gene expression^49,50^.

In Fig. 5a we can see that only 60% of the greedy path members for the SHH pathway is reported in our integrated dataset. As a matter of fact, the sequence of molecular events in this pathway is not completely understood yet. SHH signalling is critical for stem cell maintenance, embryonic development and growth^51^. It is known that the cellular response to an SHH signal is controlled by the transmembrane proteins PTCH1 and Smoothened (SMO), but the way in which SMO connects to the target zinc-finger transcription factors GLI1, GLI2 or GLI3 is still under discussion^51,52^. The geometric-based reconstruction of this pathway suggests that the proteins in charge of GLI2 activation are NEDD4 and SMAD3 (see Fig. 5e) but they are not reported pathway members. Nevertheless, we found experimental evidence for this scenario. First, Luo and colleagues measured the interaction between SMO and NEDD4 and, by means of over-expression and knock-down experiments, they identified the positive regulation of the SHH pathway by the latter^52^. Secondly, Dennler *et al.* showed that the activation of GLI2 by SMAD3 is possible *in vitro* and *in vivo*^51^. Third, there is accumulating evidence placing the NEDD4 family of E3 ubiquitin ligases as key regulators of GLI^53–55^. This information supports what the geometry of the hPIN put forward and encourages further exploration of the involvement of NEDD4 and SMAD3 in SHH signal transduction.

Finally, we examined how the perturbation of pathway members affects the routeing of signals. This is analogous to carrying out knock-out experiments, where a gene is silenced to study the effects of its absence. We observed that if *β*-catenin (CTNNB1) is removed from the hPIN, greedy routeing a signal from LRP5 to TCF7 always fails, which is expected given the dependence of TCF7 on *β*-catenin for activation^56^. However, if AXIN1 is removed, it is possible to reach TCF7 via DKK1 and LRP6 (see Fig. 5d). This alternative greedy path has to be carefully interpreted. Although it would appear that TCF7 can be activated by the binding of DKK1 to LRPs, DKK1 is known to be a negative regulator of Wnt signalling^49^. It turns out that DKK1 is one of the genes whose expression is activated by TCF7^49^. The resulting DKK1 proteins participate in a negative feedback loop that blocks Wnt signal transduction by DKK1 binding to LRPs^49,50^. This means that the LRP-DKK1-TCF7 path represents the route of an inhibitory signal rather than a stimulatory one. The combination of the undirected hPIN with a directed regulatory network, together with the inclusion of inhibitory and activating protein interaction effects, is thus key to making sense of these kinds of results^57^ and underscores the importance of interpreting geometric-based outcomes in the appropriate biological context.

## Conclusions

We used manifold learning and maximum likelihood estimation to embed the human protein interactome into the two-dimensional hyperbolic plane ^21^. Our results highlight that the latent geometry of the hPIN accurately reflects its structure and dynamics and represents a powerful tool to gain insights into the intricacies underlying this complex molecular machine.

On the one hand, the radial positioning of nodes (i.e. the geometric abstraction of their popularity or seniority status in the network) encapsulates information about the conservation and evolution of proteins. On the other,their angular positioning (abstracting the similarity between system components) captures the functional and spatial organisation of the cell. Together, the inferred radial and angular coordinates of nodes can be used to compute hyperbolic distances and assess whether two proteins are likely to interact. In addition, hyperbolic coordinates and distances can be used to simulate cell signalling events, reconstruct signal transduction pathways and study the effects of perturbations in such protein communication channels.

It is important to stress that the hPIN used throughout this article is an aggregate of protein interactions that take place under different time scales, conditions and tissues. Consequently, the results obtained by means of the latent geometry of the hPIN must be interpreted in the right biological context in order to reach sound conclusions. Notwithstanding this caveat, the use of this mapping not only reduces the universe of possibilities to test in the laboratory but can also lead to a better understanding of the mechanisms underlying the onset and development of complex human disorders. To support this endeavour, we have developed a web tool for the geometric analysis of the hPIN (http://cbdm-01.zdv.uni-mainz.de/~galanisl/gapi). With it, users can easily relate the position of proteins of interest with that of age or functional clusters and can simulate the transduction of signals between any two proteins utilising greedy routeing.

## Methods

### Protein interaction network construction

The high-quality human PIN used here represents a stringent subset of release 2.0 of the Human Integrated Protein-Protein Interaction rEference (HIPPIE)^22,23^. HIPPIE retrieves interactions between human proteins from major expert-curated databases and calculates a score for each one, reflecting its combined experimental evidence. This score is a function of the number of studies supporting the interaction, the quality of the experimental techniques used to measure it and the number of organisms in which the evolutionary relatives of the interacting human proteins interact as well. In this paper, only interactions that belong to the largest connected component were considered. To test the validity of our findings under varying levels of noise, we constructed hPINs using confidence scores ≥ {0.69, 0.70, 0.71,0.72,0.73}. The 0.72-network was preferred because it has the highest percentage of edges supported by more than one experiment. This network comprises *N* = 10, 824 nodes and *L* = 66,154 edges. Detailed structural information about all networks is listed in Table S1 and the networks themselves are provided in Supplementary Data S1. The complete HIPPIE database is available at http://cbdm-01.zdv.uni-mainz.de/˜mschaefer/hippie/download.php.

### Protein age determination

To determine the birth-time of hPIN nodes, we grouped proteins from the manually curated database SwissProt based on near full-length similarity and/or high threshold of sequence identity using FastaHerder2^58^. Briefly, age was assigned to human proteins according to the oldest common ancestor of its orthologs (sequences in different species that evolved from a common ancestor by speciation). For example, if a protein was found only in humans, it would have emerged recently, and it is considered a very young protein. If it had orthologs only in mammalian species, it would have appeared at the onset of mammals. If it had orthologs only in vertebrates, it would have emerged at the onset of vertebrates, and so on. Proteins with orthologs in all extant organisms would be the oldest. The resulting age groups, from oldest to youngest, were: 6-Cellular organisms, 5-Metazoa, 4-Chordata, 3-Mammalia, 2-Euarchontoglires and 1-Primates. The random assignment of ages (see Figs. S1b,c) was done by shuffling real ages amongst all network proteins.

### Identification of proteins classes and cancer-related proteins

We integrated information from several resources to identify proteins with transcription factor, receptor, transporter or RNA-binding activity; as well as constituents of the cytoskeleton, proteins involved in ubiquitination/proteolysis and cancer proteins. Transcription factors were identified with the aid of the Animal Transcription Factor Database 2.0^59^, the census of human transcription factors^60^ and the Human Protein Atlas^61^. The latter also helped with the identification of constituents of the cytoskeleton, proteolysis- and cancer-related proteins, receptors, transporters and RNA-binding proteins. Additional receptors and transporters were taken from the Guide to Pharmacology^62^. We also took into account RNA-binding proteins from the RNA-binding protein census^63^. The members of all these proteins classes are reported in Supplementary Data S2.

### Mapping the human protein interactome to hyperbolic space

We embedded the human protein interaction network to ℍ^2^ using LaBNE+HM^21^, an approach that combines manifold learning^24^ and maximum likelihood estimation^25^ to uncover the hidden geometry of complex networks. LaBNE+HM expects a connected network as input, typically the largest connected component or LCC. The rest of the components cannot be mapped due to the lack of adjacency information relative to the LCC. The Laplacian-based Network Embedding (LaBNE), in charge of the manifold learning part of the algorithm, generates a first geometric configuration of a network in ℍ^2^. This intermediate mapping is then passed on to HyperMap (HM), a maximum likelihood estimation method that finds node coordinates by replaying the network’s hyperbolic growth and, at each step, maximising the likelihood that it was produced by the PSM^25^. The parameter values used in the embedding process of all analysed networks are listed in Table S1. We refer the reader to^21^ for a detailed description of LaBNE+HM and its required parameters.

### Link density computation

We define link density here as the observed number of links *l* between *n* nodes, divided by the number of possible links that can occur between them, i.e. *n*(*n* − 1)/2. Since *l* varies greatly depending on the nodes being considered, we min-max normalised the link density to more easily visualise the difference between node groups. Link densities within and between age groups were compared with distributions of densities resulting from 100 random age assignments via a z-test (see Fig. 1b and S1b).

### Gene Ontology and KEGG pathway enrichment analyses

Gene Ontology^64^ and KEGG pathway^44^ enrichment analyses were carried out with RDAVIDWebService, an R interface to the Database for Annotation, Visualization and Integrated Discovery (DAVID)^65^. Only Gene Ontology terms and KEGG pathways enriched at the 0.05 significance level after Benjamini-Hochberg correction were considered.

### Clustering in the similarity dimension

We sorted the proteins increasingly by their inferred angular coordinates and computed the difference between consecutive angles with the aim to identify gaps separating groups of proteins in the similarity dimension. To determine cluster memberships, we chose the gap size *g* that produced clusters with a minimum of 10 members (*g* = 0.0132, Fig. S2). Neighbouring clusters with similar biological functions and cellular localisations were merged to avoid redundancy. Note that choosing *g* → ∞ results in a single similarity-based cluster containing all proteins, whereas *g* → 0 results in each protein forming its own cluster. The former case corresponds to analysing the function of the hPIN as a whole. The latter corresponds to analysing the role of individual proteins.

We checked if protein classes agglomerate non-randomly within their corresponding similarity-based clusters by carrying out a Fisher’s exact test. For this, we compared the proportion of proteins in class *c* = {transcription factor, receptor, transporter, RNA-binding, cytoskeleton, ubiquitination/proteolysis} that fall within a related similarity cluster against the proportion of proteins of the same class in the remaining clusters. The protein classes and their related clusters are: transcription factor, cluster 8; receptor, cluster 12; transporter, clusters 4, 5, 9 and 13; RNA-binding, clusters 7 and 14; cytoskeleton, cluster 3; ubiquitination/proteolysis, clusters 1, 2 and 15. The resulting p-values were adjusted with the Benjamini-Hochberg method.

### Protein interaction prediction

The hPIN can be represented by an undirected graph *G*(*V*,*E*), where *E* is the set of interactions between a set *V* of proteins. When link prediction methods are applied to a network, they assign likelihood scores of interaction to all the disconnected node pairs. We ranked these candidate protein interactions by hyperbolic distance and compared the 100 closest disconnected protein pairs with the top-100 predictions produced by different classes of prediction methods: the neighbourhood-based link predictors Common Neighbours (CN)^66^, Adamic & Adar (AA)^67^ and Preferential Attachment (PA)^66^; the Cannistraci-Alanis-Ravasi index (CAR) and the CAR-based AA (CAA) and PA (CPA)^68^; the embedding-based link predictors ISOMAP^9,10,30^and non-centred Minimum Curvilinear Embedding (ncMCE)^14^; and the recently proposed Structural Perturbation Method (SPM)^33^. Neighbourhood-based predictors assign high likelihood scores to node pairs that share many neighbours. For CAR-based predictors, these common neighbours have to connect with each other. Embedding-based methods rank non-adjacent nodes through distances on a network projection in two-dimensional Euclidean space. Finally, SPM hinges on the hypothesis that links are predictable if removing them has only small effects on network structure. We refer the reader to^32,33^for more details and predictor formulations.

The discrimination between good and bad candidates was based on the Guilt-by-association Principle, which states that if two proteins are involved in similar biological processes or are located in the same cellular compartment, they are more likely to interact^69^. Thus, good candidate interactions correspond to top-ranked pairs of proteins that play a role in at least one common pathway (functional homogeneity) or locate to the same subcellular structure (localisation coherence). This link prediction evaluation framework is extensively used for biological networks^24,70–73^. Pathway memberships were determined via KEGG pathways^44^ and cellular localisations via the Cellular Compartment aspect of the Gene Ontology^64^ and the Cell Atlas^74^. The top-100 candidate interactions of each link predictor are provided in Supplementary Data S6.

### Greedy routeing and pathway reconstruction

Greedy routeing of signals involved 100 experiments with 1000 sources and 1000 targets each. Sources and targets were selected at random or from a pool of transcription factors, receptors or cancer-related proteins. Since the number of proteins in each one of these classes differs, the pools were formed by 500 randomly-selected members of each one. Mann-Whitney U tests were used to compare the distribution of efficiencies achieved in the ℍ^2^ case with the distribution from the Rec-TF case (see Fig. 4a).

For pathway reconstruction, we computed greedy and shortest paths from one or more sources to one or more targets from the 24 signal transduction pathways listed by the Kyoto Encyclopedia of Genes and Genomes (KEGG)^44^ and their equivalents in Reactome^45^ and WikiPathways^46^. These starting- and end-points were determined based on KEGG itself and the literature^37,38^and represent canonical transduction initiators and transcriptional regulators, respectively. For a pathway *P_i_* with *m* members *P_i_* = {*p_1_,p_2_, …, p_m_*}, we simulated the transduction of a signal from *s* ∈ *P_i_* to *t* ∈ *P_i_* by calculation of the greedy or shortest path between *s* and *t*. Then, we computed the fraction of proteins that form the greedy or shortest path and that are members of *P_i_*. For some pathways, we compiled more than one *s* — *t* pair and computed the average fraction instead. All these pairs and their corresponding pathways are reported in Supplementary Data S7. Pathway membership was determined by integrating data from KEGG, Reactome and WikiPathways (results reported in Fig. 5a and Fig. S10a) or by using each resource separately (results reported in Fig. S10b-d).

### Hardware used for experiments

All the experiments presented in this paper were executed on a Lenovo ThinkPad 64-bit with 7.7 GB of RAM and an Intel Core i7-4600U CPU @ 2.10 GHz × 4, running Ubuntu 16.04 LTS. The only exceptions were the signal delivery, connection probability and link prediction experiments, which were executed on nodes with 64 GB of RAM, within the Mogon computer cluster of the Johannes Gutenberg Universität and the computer cluster of the Institute of Molecular Biology.

## Data availability

All relevant used and produced data accompanies this text as Supplementary Information:

- **Supplementary File.** Contains Supplementary Table S1 and Figures S1-S10.
- **Supplementary Data S1.** The five different human protein interaction networks (hPINs).
- **Supplementary Data S2.** Inferred hyperbolic coordinates, ages and classes of the proteins in the five constructed hPINs.
- **Supplementary Data S3.** GO and KEGG pathway enrichment analyses of age groups from the 0.72- hPIN.
- **Supplementary Data S4.** GO and KEGG pathway enrichment analyses of similarity-based clusters from the main 0.72-hPIN.
- **Supplementary Data S5.** GO and KEGG pathway enrichment analyses of Louvain communities from the main 0.72-hPIN.
- **Supplementary Data S6.** Top-100 protein interactions predicted by ten different link predictors.
- **Supplementary Data S7.** Considered signal transduction pathways, sources and targets.

## Acknowledgements

The authors gratefully acknowledge the computing time granted on the supercomputer Mogon at the Johannes Gutenberg Universität and the computer cluster at the Institute of Molecular Biology in Mainz.

## Competing interests

The authors declare that they have no competing interests.

## Author’s contributions

GAL created LaBNE+HM, designed, implemented and carried out the experiments. GAL and PM performed the data integration. PM was in charge of the protein age assignment and of receptor and transcription factor identification. MAAN supervised the research. GAL wrote the manuscript, incorporating comments, contributions and corrections from PM and MAAN. All authors read and approved the final version of the manuscript.

